# Using a two-stage convolutional neural network to rapidly identify tiny herbivorous beetles in the field

**DOI:** 10.1101/2021.05.27.445368

**Authors:** Hironori Takimoto, Yasuhiro Sato, Atsushi J. Nagano, Kentaro K. Shimizu, Akihiro Kanagawa

## Abstract

Recently, deep convolutional neural networks (CNN) have been adopted to help beginners identify insect species from field images. However, the application of these methods on the identification of tiny congeneric species moving across heterogeneous background remains difficult. To enable rapid and automatic identification in the field, we customized a method involving real-time object detection of two *Phyllotreta* beetles. We first performed data augmentation using transformations, syntheses, and random erasing of the original images. We then proposed a two-stage method for the detection and identification of small insects based on CNN, where YOLOv4 and EfficientNet were used as a region proposal network and a re-identification method, respectively. Evaluation of the model revealed that one-step object detection by YOLOv4 alone was not precise (Precision = 0.55) when classifying two species of flea beetles and background objects. In contrast, the two-step CNNs improved the precision (Precision = 0.89) with moderate accuracy (F-measure = 0.55) and acceptable speed (ca. 5 frames per second for full HD images) of detection and identification of insect species in the field. Although real-time identification of tiny insects remains a challenge in the field, our method aids in improving small object detection on a heterogeneous background.

## Introduction

Insects are some of the most abundant groups of animals in the terrestrial ecosystem. Rapid and accurate identification of insect species is essential for pest management (Park et al., 2020; Preti et al., 2020) and species-level identification in biodiversity monitoring (Hogeweg et al., 2019; Høye et al., 2021). To augment the expertise in insect taxonomy and ecology, several attempts have been made to develop automated identification systems (Martineau et al., 2017). However, handcrafted extraction of image features requires experience on image processing (Mayo and Watson, 2007; Samanta and Ghosh, 2012; Yalcin, 2015; Xie et al., 2015; Rani and Amsini, 2016). Recently, the deep convolutional neural network (CNN) has revolutionized this task by facilitating the automatic extraction of key features from a huge number of insect images (Valan et al., 2019; Park et al., 2020; Almryad and Kutucu, 2020; Hansen et al., 2020; Høye et al., 2021).

Despite its promising implications, the usage of CNN is largely restricted to a snapshot image taken on a simple background, such as a capturing sheet (Hansen et al., 2020) and trapping system (Preti et al., 2020). In contrast, field images are much more complex since they involve a heterogeneous background with multiple species and individuals (Martineau et al., 2017). Furthermore, a snapshot or interval camera may not be sufficient to capture the key characteristics of species identification (Preti et al., 2020). A series of images, namely a video, may solve this issue by capturing multiple posing for an individual insect (Høye et al., 2021). Thus, a field video may provide a unique material for a quick and thorough imaging of a target insect.

In the context of computer vision, CNN-based object detection has now enabled fast and accurate processing of a large series of images that involve living organisms and artificial objects (Liu et al., 2019). The goal of object detection is not only to determine whether objects from general categories are present on an image, but also to return the spatial location and determine the extent of each object by using a bounding box. A CNN is designed to automatically learn the spatial-feature hierarchies layer by layer and the consequent sub-sampling layers in a feature hierarchy that results in an inherent multiscale pyramid (Yamashita et al., 2018). It is well recognized that the higher layers are the most robust in terms of object pose, illumination, part deformation, and other variations because they have large receptive fields and strong semantics. However, a critical disadvantage in this step is the loss of geometric details and the low resolution, such that small objects more likely lack critical features since they are processed in subsequent layers (Liu et al., 2019). Thus, the detection of small objects using general-purpose CNNs is challenging.

In general, a large training dataset is required for CNN models (LeCun et al., 2015). However, currently available insect images in public datasets may not be sufficient to train CNN models for real-world variations in object angles, camera conditions, and field background. To efficiently train CNN models for an individual insect, we need to take multi-angle photographs in a systematic manner (Hansen et al., 2020; Høye et al., 2021). However, in this method, the background is likely homogenous, which is not the case in fields with complex and heterogenous background. In contrast, citizen scientists may provide a large number of field photographs (Silvertown et al., 2015; Van Horn et al., 2018), but the database may be biased with respect to species, angles, and background. Synthesis or transformation of original images, called data augmentation (Shorten and Khoshgoftaar, 2019), is thus required to create effective variation in training data.

The objectives of this study were (i) to test whether an existing method of real-time object detection could be used in a field video of small insects and (ii) to improve the detection of small insects in the field. Specifically, we proposed a two-stage detection and identification method for small insects based on CNN (Fig. 1). For real-time insect detection, we focused on an object detection method called You Only Look Once (YOLO) v4 (Bochkovskiy et al., 2020). YOLOv4 is a single-shot object detector that is trained end to end from raw pixels to ultimate categories. When the size of a target object is relatively large, YOLOv4 can detect the object accurately (Ovchinnikova et al., 2021). Contrastingly, when the size of the target object is small, the object is often undetected or over-detected by YOLOv4. To address this issue, we first used YOLOv4 as a region proposal network that detects many possible candidate regions. All detected region proposals were then re-identified by a CNN-based classifier, the EfficientNet (Tan and Le, 2019). With a less computational cost than other models, the EfficientNet retains sufficient capacity over various benchmark datasets by adopting a compound scaling method that enlarges the network depth, width, and resolution. In addition, we proposed a data augmentation method to train the two-stage CNNs. Our proposed method aimed to address specific problems regarding the size, appearance, and training data of insects in the field.

**Figure 1:**
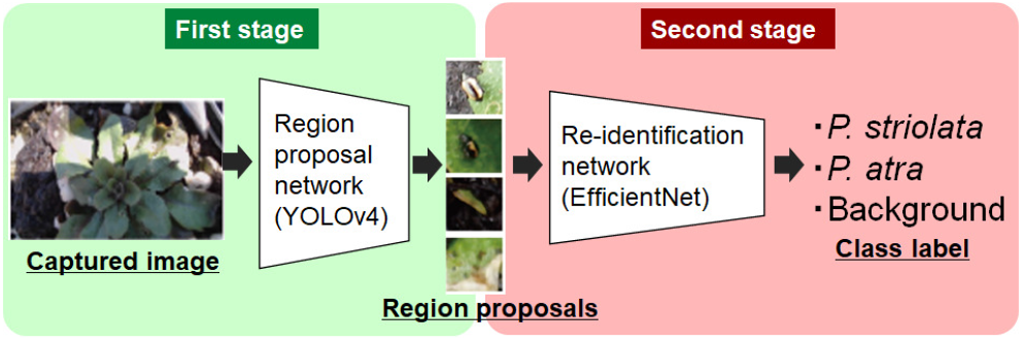
Flow of the proposed two-stage detection method. During the first stage (left), the proposed method detected several region proposals from an input image by YOLOv4 with a low threshold of detection. In the second stage (right), all region proposals were re-identified using the EfficientNet.

## Material and methods

### Study species and site

The target insects were the striped flea beetle *Phyllotreta striolata* and the turnip flea beetle *Phyllotreta atra*. The two insect species were also known as pests of *Brassica* crops (Ahuja et al., 2010). These insect species often appeared on the genus *Arabidopsis* (Brassicaceae) during the early summer days (Sato et al., 2019; Shimizu-Inatsugi et al., 2021). In this study, a total of 3800 potted individuals of *Arabidopsis thaliana* were transplanted in the field garden at the University of Zurich at the Irchel campus (Zurich, Switzerland; 35°06′N, 134°56′E) in July 2017 (1600 pots), July 2018 (1600 pots), and July 2019 (600 pots). Fig. 2 shows a sample image of the two insect species appearing on an *A. thaliana* plant. Both species had a length of approximately 2 mm and had similar habitus. The only difference in their appearance was the stripes on their back. *P. striolata* had yellow stripes, while *P. atra* had none. Furthermore, adult beetles created small holes when eating plant leaves, which might explain the appearance of gaps between the leaves (a lower blue square of Fig. 2) or a leaf hole in the backgrounds of images. The formation of these holes might be attributed to the feeding behavior of flea beetles (an upper blue square of Fig. 2).

**Figure 2:**
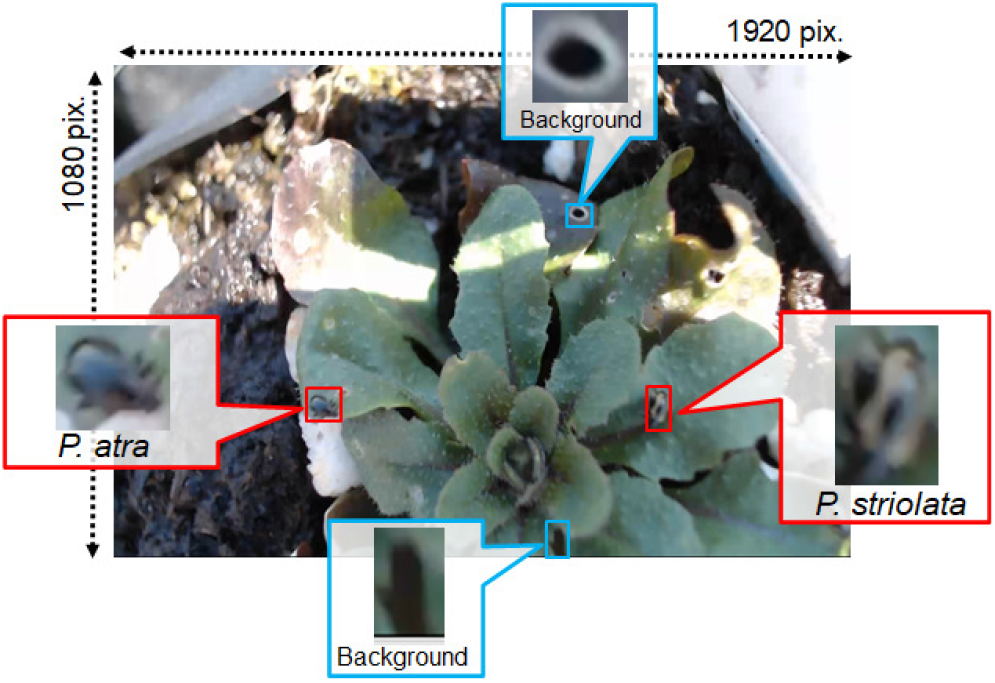
Example of a captured image in a real field environment. Two species of *Phyllotreta* beetles (red square; ca. 2 mm in body length) simultaneously appeared on an individual *Arabidopsis thaliana*.

### Data collection

For training data, we collected images of *P. striolata* and *P. atra* via Google search. Our own photographs were taken during the transplant experiment in 2017 and 2018. We used a RICOH WG-4 digital camera (RICOH Company, Ltd., Toky Japan), which generated 258 photographs (1,920×1,080 pixels) for *P. striolata* only, 106 photographs for and *P. atra* only, and 41 photographs of both *P. striolata* and *P. atra*. For the Google search, we used the following keywords: “*Phyllotreta striolata*”, “*Phyllotreta atra*”, “striped flea beetle” (common name of *P. striolata* in English), “turnip flea beetle” (common name of *P. atra* in English), “kisuji-nomi-hamushi (common name of *P. striolata* in Japanese, searched in Japanese katakana letters), or “mumon-kisuji-nomi-hamushi” (common name of *P. atra* in Japanese, searched in Japanese katakana letters). Hence, our search resulted in a total of 142 and 49 images for *P. striolata* and *P. atra*, respectively. However, due to the low resolution of the images and the high similarity of the two species, the effective number of high-resolution images with different posing for training data was only 32 and 29 for *P. striolata* and *P. atra*, respectively. The original photographs taken during our own fieldworks were deposited on the GitHub repository (https://github.com/h-taki/identification-of-tiny-insects).

To procure a test video, we used a full HD video mode (1,920×1,080 pixels) of Logicool C920r camera (Logicool Co. Ltd., Lausanne, Switzerland). A test video was photographed in 2019 to differentiate the survey years between the training and test data. The camera was positioned sufficiently close to but somewhat far from the target insects to let flea beetles escape. To set a ground truth, we annotated the two species on every several frames of an entire video, which resulted in the generation of 215 annotated video frames. The series of annotated frames captured 646 insect objects, with 427 (*P. striolata* and 219 *P. atra*. The annotation was made using the LabelImg tool (https://github.com/tzutalin/labelImg). The annotated test video is available as Supplementary Video S1.

### Data augmentation

Although optimizing many parameters of CNN based on deep learning required a large dataset, there were only small datasets for pest detection and identification. Existing datasets only had approximately 100 images for each type (Samanta and Ghosh, 2012; Rustia et al., 2018; Xia et al., 2018; He et al., 2019; Wang et al., 2012; Venugoban and Ramanan, 2014; Xie et al., 2018; Deng et al., 2018). To obtain a more generalized model, training data should have fine diversity as the objects varied in size, lighting conditions, and poses. To circumvent the limited quantity and diversity of training data, data augmentation was a useful technique that could increase the size of the training set without acquiring new images. Data augmentation was typically performed as part of dataset preprocessing for training an image detection and classification model, such as the CNN-based model. The procedure included cropping, flipping, scaling, rotating, adding noise, and color transformation.

In this study, we used augmented datasets for the two-stage detection. First, we proposed data augmentation to effectively train YOLOv4 for the sake of a region proposal network. Real segmented objects (i.e., segmented insect images) were pasted into natural images, such as photographs of soils and/or plants. Examples of synthesized images are shown in Fig. 3. A mask image as a foreground image was created by extracting only insect regions from the original images. A single image was then randomly selected from a large natural image dataset that did not contain any insects. A background image was cropped at 512×512 pixels from the selected image. The procedure of data argumentation consisted of conventional image processing methods, image synthesis (Fig. 3c), and random erasing (Fig. 3a, b). First, the foreground and background images were subjected to image processing operations such as cropping, flipping, and scaling. By combining a foreground image with a background image, we constructed a large training image dataset. Overlapping foreground images were allowed when combining multiple foreground images into a single background image. After the foreground and background images were combined, additional image processing operations were conducted using the Gaussian noise, gamma transform, and Gaussian to further diversify the synthesized images (e.g., Fig. 3c). In addition to these classical methods of data augmentation, we finally applied random erasing methods to the dataset (e.g., Fig. 3a, b). Random erasing generated training images with various levels of occlusion by randomly selecting a rectangle region in an image whose pixels were erased with random values (Zhong et al., 2020). In a captured image in a field environment, insects were partially hidden by self-occlusion, plant leaves, and soil particles. By conducting random erasing to insect-specific issues, we reduced the risk of network overfitting and made the model robust to occlusion. For the data augmentation for training YOLOv4, the parameters are shown in Table 1a.

**Figure 3:**
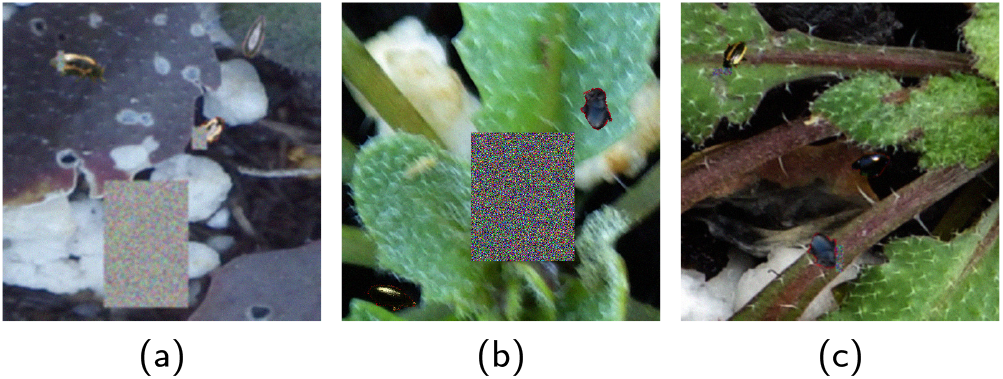
Examples of data augmentation for the first stage using the YOLOv4. Training data were synthesized using foreground pictures of flea beetles and background field images. The squares in the panels (a) and (b) indicated a randomly erased region.

**Table 1.** The parameters of the data augmentation for training YOLOv4 (a) or EfficientNet (b).

| (a) For YOLOv4 |  |
| --- | --- |
| Size of the foreground object | 40 - 70 pixels |
| Maximum number of foreground object contained in one image | 3 |
| Scaling rate (while ignoring aspect ratio) | 80 - 120% |
| Gaussian noise $\sigma$ | 0.0 - 9.0 |
| Gamma transform $\gamma$ | 0.5 - 1.5 |
| Gaussian blur, $\sigma$ | 1.0 - 3.0 |
| Gaussian blur, filter size | 3 - 11 |
| Erasing area ratio | 2 - 30% |

| (b) For EfficientNet |  |
| --- | --- |
| Size of the foreground object | (B0) 149 - 187 pixels,<br>(B1) 160 - 200 pixels,<br>(B2) 173 - 217 pixels,<br>(B3) 200 - 250 pixels |
| Maximum number of foreground object contained in one image | 1 |
| Scaling rate (while ignoring aspect ratio) | 80 - 120% |
| Gaussian noise $\sigma$ | 0.0 - 9.0 |
| Gamma transform $\gamma$ | 0.5 - 1.5 |
| Gaussian blur, $\sigma$ | 1.0 - 3.0 |
| Gaussian blur, filter size | 3 - 11 |
| Erasing area ratio | 2 - 40% |

Second, we proposed data augmentation to train the EfficientNet as a classifier. The basic concept of this procedure was the same as that in the data extension for YOLO, where we combined a masked foreground image with a background image that did not contain any insects. At most, one foreground image was composited. For the foreground, background, and combined images, the same image processing as the data augmentation for YOLOv4 was applied. For the data augmentation employed to train the EfficientNet, the parameters are shown in Table 1b. Fig. 4 shows an example of data augmentation for the second stage of the proposed augmentation method.

**Figure 4:**
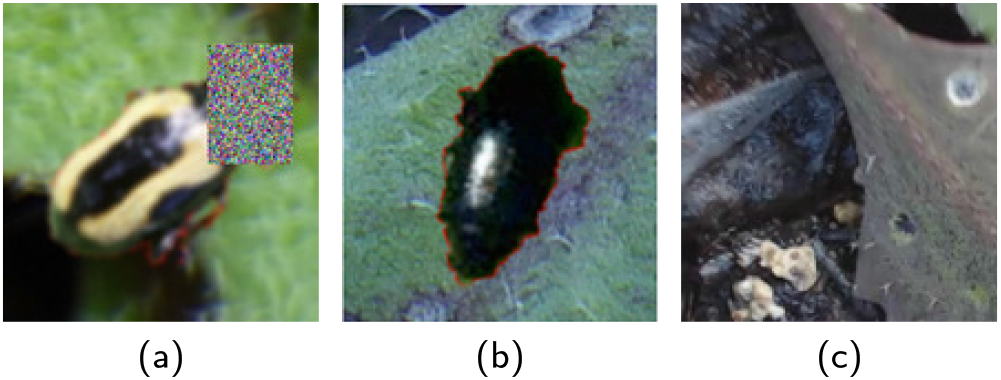
Examples of data augmentation for the second stage using the EfficientNet. (a) A foreground image of *P. striolata* was synthesized on a background image with random erasing; (b) A foreground image of *P. atra* was synthesized on a background image; (c) A background image in which only the image processing method (Table 1b) was applied.

### Insect candidate detection using a CNN-based object detector

In the emerging area of deep learning, effective and efficient CNN-based object detection models have been proposed so far, including faster R-CNN (Ren et al., 2017), Region-based Fully Convolutional Networks (R-FCN) (Dai et al., 2016), Single Shot multibox Detector (SSD) (Liu et al., 2016), and YOLO(Redmon et al., 2016). Two-stage detectors such as faster R-CNN and R-FCN had a relatively high accuracy of detection but unrealistic computational costs. By contrast, one-stage detectors such as YOLO and SSD had a lower accuracy but had an increased speed.

Redmon et al. (2016) proposed YOLO for the unified detector casting object detection as a regression problem from image pixels to spatially separated bounding boxes and associated class probabilities. Unlike region-based approaches, YOLO lacked the region proposal generation stage. Therefore, YOLO directly detected objects using a small set of candidate regions. In particular, YOLO divided an image into an *S* × *S* grid, from which *C* class probabilities, *B* bounding box locations, and confidence scores were predicted.

Bochkovskiy et al. (2020) created YOLOv4, which was an improved version of YOLO. In YOLOv4, the CNN-based feature extractor was replaced by the CSPDarknet53 feature extractor. YOLOv4 had achieved excellent scores on standard detection tasks and balanced speed and accuracy well. However, YOLOv4 might fail in detecting small objects, plausibly because of the coarse grid division and because each grid cell could only contain a few objects.

Given the disadvantage of YOLO in detecting small objects, we used YOLOv4 as a region proposal network during the first stage of the proposed method. For each image, YOLOv4 generated a set of bounding boxes with their confidence scores ranging from 0 to 1. Confidence score pertained to the level of confidence on whether the box contained the object. To thoroughly screen insect regions, we set the threshold of confidence score to a low value of 0.2. This led to the overdetection of background objects and insects. In comparison, the threshold of confidence score for an object detection task by the YOLO series was often set to a value that ranged from 0.3 to 0.5 (Padilla et al., 2021; Ovchinnikova et al., 2021; Redmon et al., 2016; Redmon and Farhadi, 2018).

### Insect identification using a CNN-based classifier

In the second stage of the proposed method (Fig. 1), the region proposals detected by YOLOv4 were re-identified as insects or not. We employed the EfficientNet as a classifier during the second stage.

The past development of CNN was synonymous to the increased depth of the network. By increasing the depth and widening the channel size, a more complex network achieved a high resolution of the image data. In addition, it resulted in more fine-grained characteristics. This development in the network size improved the classification accuracy of the network, but also led to the problem of the high computational cost of the gradient explosion parameter. ResNet (He et al., 2015) proposed that skip connection could avoid gradient explosion skillfully. MobileNet (Howard et al., 2017) used pointwise and depthwise convolutions to reduce network parameters and improve training efficiency. The SENet (Hu et al., 2017) weighted various features by loss of network training in order to achieve better results in model training.

The EfficientNet integrated characteristics of the above networks through uniformly scaling all dimensions of depth-width resolution using a compound coefficient. Unlike conventional practices that arbitrarily scaled these factors, the EfficientNet scaling method uniformly scaled the network width, depth, and resolution with a set of fixed scaling coefficients. The formula for calculating the composite proportion coefficient was as follows.

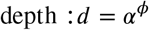

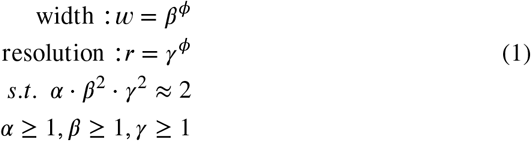

where *α*, *β*, *γ* were constant coefficients determined by a small grid search on the original small model. *ϕ* was a user-defined coefficient that controlled the number of resources that were available for model scaling. The compound scaling method was justified by the intuition that if the input image was bigger, more layers and channels were needed by the network to increase the receptive field and to capture more fine-grained patterns on the bigger image, respectively. There were eight types of the EfficientNet, from the EfficientNet-B0 to the EfficientNet-B7, with an increasing network scale.

Because the size of the region proposal depended on the output of YOLOv4, an input image for the EfficientNet was resized to the required size of the EfficientNet irrespective of the aspect ratio. To re-identify the insects with an over-detection of the suppressed background region, the proposed EfficientNet had three corresponding output classes: *P. striolata*, *P. atra*, and the background. The Softmax function was used for the output layer of the re-identification task that posed a multi-class classification problem. A standard Soft-max cross-entropy was used as a loss function.

### Model evaluation

To demonstrate the effectiveness of the proposed two-stage method for the insect detection, we first applied YOLOv4 alone for detection, and then compared the results of the first stage with those of the two-stage method. For data augmentation, we evaluated the effectiveness of random erasing.

We adopted a sliding window approach to input the full captured frames into YOLOv4. We chose 512×512 pixels as an input image size for YOLO. When the image was resized to 512×512 pixels (where the frame size of the original captured image was 1,920×1,080 pixels), insects that were 2 mm in length were too compacted that even human experts could not identify them. To solve this issue, we cut an original captured image into eight pieces (800×800 pixels with overlap allowed). We then inputted them into the YOLOv4 (Fig. 5). Each cropped image was then resized to 512×512 pixels to fit within the required input size for YOLOv4.

**Figure 5:**
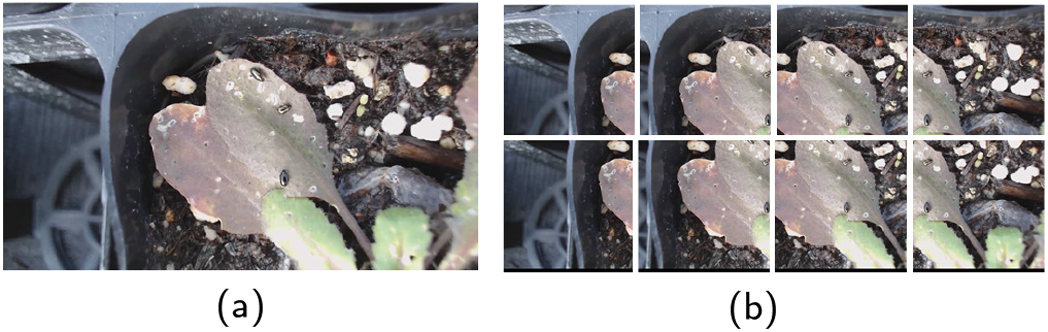
Sliding window approach was used to input data to the YOLOv4. An original image (1,920×1,080 pixels) was split into eight images (800×800 pixels), and overlap was allowed. The images were then resized to 512×512 pixels for input to YOLOv4.

To train the YOLOv4, we prepared 12,000 images using the data augmentation method proposed above. Out of all the augmented images, 80% of the images were used for training, while the remaining 20% was for validation. The YOLOv4 was trained in 150 epochs. The batch size was set at 64.

To re-identify the region proposal, we inputted outputs from the YOLOv4 into the EfficientNet. As mentioned above, there were eight types of the EfficientNet. Starting from the baseline EfficientNet-B0, the baseline network was scaled up from EfficientNet-B1 to B7 based on the compound scaling method with different *ϕ* values. The EfficientNet-B0 had the smallest network size, as it had an input image resolution of 224×224. The size of the region proposal detected by the YOLOv4 roughly ranged from 40×40 to 150×150 pixels; therefore, all detected region proposals were resized to the input size of the EfficientNet regardless of the aspect ratio. When the target image was small, re-identification using a large network did not improve the accuracy but merely increased the computational cost. In this study, we chose four relatively small-sized networks (i.e., EfficientNet-B0, -B1, -B2, -B3) as a classifier. We also compared the four sizes of the EfficientNet with another classifier, Xception (Chollet, 2016). Xception (meaning “extreme inception”) was an extension of the inception architecture that replaced the standard inception modules with depth-wise separable convolutions.

To train the EfficientNet, we prepared 4,000 images for each category using the data augmentation method proposed above. We used 80% of the 4,000 images for training and the remaining 20% for validation. The EfficientNet was trained by the Adagrad in 100 epochs. The dropout rate for each task was set at 0.5. The batch size was set at 20.

All evaluations were performed on the NVIDIA GeForce RTX 2080Ti GPU with 11G memory, and the software was Windows 10, Python 3.7, CUDA 10.2. Keras 2.4.3 were used to build all models.

To compare model results with the test video, we used the Intersection over Union (IoU) as the threshold for positives or negatives based on the object confidence scores estimated by the object detector. IoU was defined as the intersection over the union of the two bounding boxes, which were the ground-truth bounding box and the predicted bounding box. In this paper, we set the threshold of IoU at 0.3 to consider correct or incorrect detection results.

## Results

First, we verified the effectiveness of random erasing to improve the performance of the YOLOv4. Tables 2 displays the confusion matrices of the one-stage detection using only the YOLOv4 with/without random erasing. In the set of tables, the rows and columns of the confusion matrix correspond to the true classes and predicted classes, respectively. The threshold of confidence score in YOLOv4 was set at 0.2 to accept the ground-truth objects predicted by the detection task. To quantify the detection performance, we calculated the precision, recall, and F-measure (Table 3).

**Table 2.** Confusion matrices of the one-stage detection using only the YOLOv4 with (a) or without (b) random erasing.

| (a) With random erasing |  |  |  |
| --- | --- | --- | --- |
|  | *P. striolata* | *P. atra* | Background |
| *P. striolata* | 171 | 4 | 0 |
| *P. atra* | 0 | 98 | 0 |
| Background | 93 | 123 | 0 |

| (b) Without random erasing |  |  |  |
| --- | --- | --- | --- |
|  | *P. striolata* | *P. atra* | Background |
| *P. striolata* | 159 | 13 | 0 |
| *P. atra* | 4 | 102 | 0 |
| Background | 115 | 138 | 0 |

**Table 3.** Detection results using only the YOLOv4 with/without Random Erasing (R. E.).

|  | Precision | Recall | F-measure |
| --- | --- | --- | --- |
| YOLOv4 w/ R. E. | 0.55 | 0.42 | 0.47 |
| YOLOv4 w/o R. E. | 0.49 | 0.40 | 0.44 |

The results showed that the detection performance was improved by subjecting the images to random erasing of the training dataset of the YOLOv4 (Table 3). By applying the YOLOv4 with random erasing to the annotated test video, we obtained 489 candidate insect regions from the YOLOv4, but 44.2% of the output was considered over-detection. This result was reasonable because the threshold of confidence score was relatively low for detecting as many as possible small insects. If over-detection was unfavorable, it could be suppressed by setting a relatively high threshold. However, the undetected rate would be even worse than the current undetected rate of 57.7%.

Table 4 shows the results in which outputs from the YOLOv4 with random erasing were re-identified using the Efficient-Net. In these results, all classifier models were trained on the dataset with random elimination applied. The results suggest that the most accurate of the proposed re-identification models were the YOLOv4 and EfficientNet-B1.

**Table 4.** Comparison of performance among the EfficientNet and Xception models.

|  | Precision | Recall | F-measure |
| --- | --- | --- | --- |
| EfficientNet-B0 | 0.86 | 0.38 | 0.53 |
| EfficientNet-B1 | 0.89 | 0.40 | 0.55 |
| EfficientNet-B2 | 0.89 | 0.39 | 0.54 |
| EfficientNet-B3 | 0.91 | 0.39 | 0.54 |
| Xception | 0.84 | 0.35 | 0.49 |

To show the effects of random erasing on the performance of the EfficientNet, Tables 5 displays the confusion matrices of the two-stage detection using the YOLOv4 with random erasing and the EfficientNet-B1 with or without random erasing. Table 6 shows the detection performance of these methods. By applying random erasing to the training dataset of the EfficientNet, performance was also improved. In addition, our proposed two-stage detection outperformed one-stage detection using only the YOLOv4.

**Table 5.**
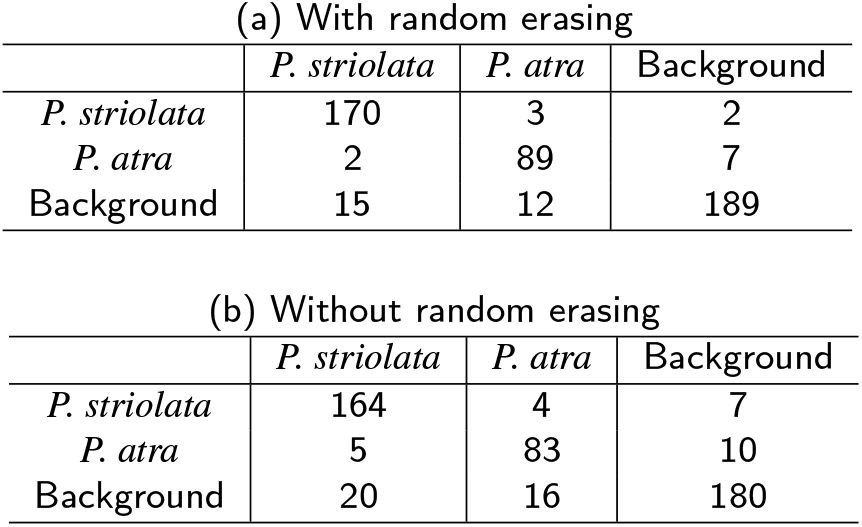
Confusion matrices of the two-stage detection using the EfficientNet-B1 with (a) or without (b) random erasing.

| (a) With random erasing |  |  |  |
| --- | --- | --- | --- |
|  | *P. striolata* | *P. atra* | Background |
| *P. striolata* | 170 | 3 | 2 |
| *P. atra* | 2 | 89 | 7 |
| Background | 15 | 12 | 189 |

| (b) Without random erasing |  |  |  |
| --- | --- | --- | --- |
|  | *P. striolata* | *P. atra* | Background |
| *P. striolata* | 164 | 4 | 7 |
| *P. atra* | 5 | 83 | 10 |
| Background | 20 | 16 | 180 |

**Table 6.** Detection results the two-stage detection with/without Random Erasing (R. E.).

|  | Precision | Recall | F-measure |
| --- | --- | --- | --- |
| EfficientNet-B1 w/ R. E. | 0.89 | 0.40 | 0.55 |
| EfficientNet-B1 w/o R. E. | 0.85 | 0.38 | 0.53 |

After re-identification by the EfficientNet-B1, 87.5% of falsely detected background areas were corrected as background classes (Supplementary Video S2). Thus, the two-stage detection greatly improved the precision compared with one-stage detection using only the YOLOv4. We also confirmed that the precision and resultant F-value were improved, while the recall was nearly the same (Table 3 and 6).

In addition, we evaluated the speed to detect and re-identify a region proposal. Table 7 shows the processing time of re-identification using each classifier model for one region proposal. In this table, “Detection with YOLOv4 and reidentification” corresponded to the frame rate, which was the frame per second (FPS) of the detection process for one frame and the re-identification process for two region proposals since the number of candidate regions detected from a single frame in this evaluation was around two.

**Table 7.** Comparison of the processing time among the re-identification models in seconds (s) and frames per second (FPS).

| Re-identification models | Classification of one image [s] | Detection with YOLOv4 and re-identification [FPS] |
| --- | --- | --- |
| EfficientNet-B0 | 0.027 | 5.4 |
| EfficientNet-B1 | 0.029 | 5.3 |
| EfficientNet-B2 | 0.030 | 5.3 |
| EfficientNet-B3 | 0.033 | 5.1 |
| Xception | 0.031 | 5.2 |

## Discussion

The paradigm of visual recognition has changed rapidly after the emergence of the ImageNet dataset, which has demonstrated the power of data-driven feature learning. During the past years, CNN has become the most effective architecture for visual recognition tasks. Therefore, many researches have been conducted on deep learning-based approaches for insect detection and identification (Lim et al., 2017; Rustia et al., 2018; Shen et al., 2018; Xia et al., 2018; He et al., 2019). These conventional methods have some disadvantages in terms of practicality, where insects can be detected only on a very simple background, such as a supplementary sheet or trap. To achieve a system that is more suitable for field use and more user-friendly, we advocated the usage of a unique application of CNN-based object detection to a video collected in the field. The present study provided a two-stage method that could greatly improve the precision to distinguish small congeners and background objects, while there remained major challenges in real field environments regarding the size, resemblance, and training data of insects.

Given the advantage of YOLO in rapid processing, we aimed to create a balance between the speed and accuracy in our two-stage method. The processing of the proposed two-stage method is approximately 5 FPS in total (Table 7), which suggests an acceptable speed in an interactive user interface. As if unfamiliar languages are translated using a smartphone camera, we suppose that users leverage the proposed system to quickly check the location and species of insects in the field. Furthermore, sedentary insects do not suddenly move out of the view of a camera. Unless the target is a fast-moving species such as an adult butterfly flying across the sky, the processing cost of the proposed method is unlikely to be a hindrance for practical use. While the current performance (namely 5 FPS) is not so smooth that human recognizes the output as a real-time video, it would be faster due to technological advances in the future.

One of the most serious issues is the difficulty in distinguishing small insects and background objects. Even though the YOLOv4 alone could distinguish two species of flea beetles, we observed over-detection simultaneously (Table 2); thus, it was plausible to use an additional CNN-based classifier to suppress over-detection. Combined with the latest object detection methods, the proposed two-stage method correctly re-identified most of the over-detections as background (Table 5). Figure 6 shows an example of correct identification by re-identification, where background objects (Fig. 6a upper) or a leaf hole (Fig. 6a lower) are initially mistaken as flea beetles but are then re-identified as background. Notably, the smallest individual that could be detected by the proposed method was 35×50 pixels in the HD size image (Fig. 7). The area ratio of the detected insect to frames was merely *<*0.1%. These results suggest that our two-stage method has a potential to distinguish small targets from various background objects.

**Figure 6:**
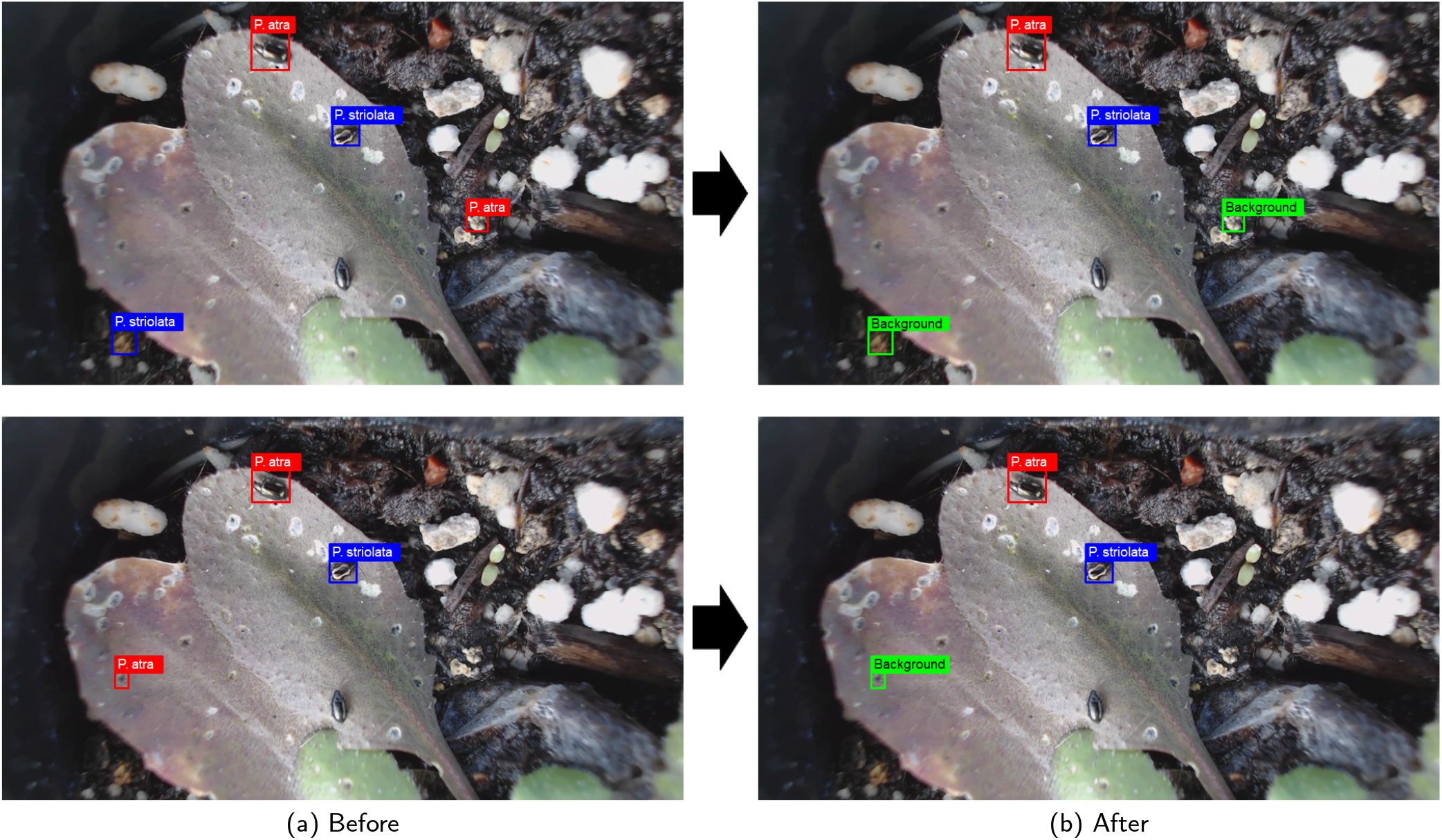
Figure 6. Example of correct identification by re-identification. Results of the first stage using the YOLOv4 (a), and results of the second stage using the EfficientNet (b). The video results of the two-stage method are available as a supplementary material (Supplementary Video S2).

**Figure 7:**
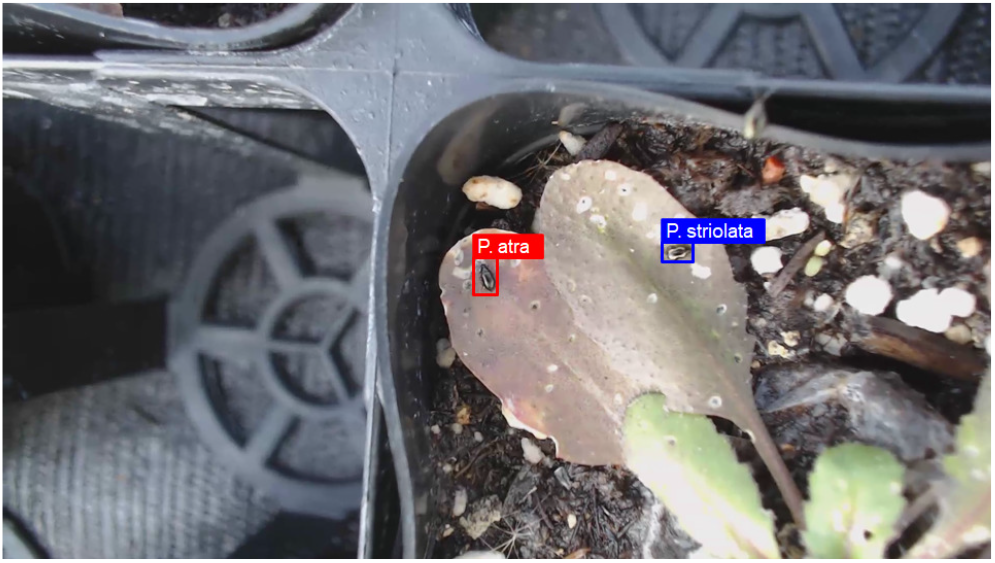
Examples of the smallest individuals that were detected correctly. The size of the original image is 1,920×1,080 pixels, where the regions of flea beetles detected are ca. 35×50 pixels. The video results are available as a supplementary material (Supplementary Video S3).

Another issue in the field is heterogeneous photographing conditions due to lighting and angles. Specifically, Figure 6 shows that a *P. atra* individual located in the lower center cannot be detected until it moves (until the 30 s mark in Supplementary Video S3). This missed detection might have occurred due to the strong light source, which was the sunlight, was reflected on *P. atra*’s black back with a small contrast between the target i.e., the black beetle and the background, which was consisted of dark-colored leaves. A high precision with a small Recall (Table 6) indicates a high identifiability with non-exhaustive searching; however, this performance is still useful for screening with an interactive user interface. In fact, even the undetected *P. atra* individual was finally detected when its appearance changed among different frames (from the 31 s mark in Supplementary Video S3). Since insects moved spontaneously, they exhibited various appearances in different times. It was also possible that movements of a camera device, including camera shaking and resultant re-focusing, led to slight changes in insect appearance. While changes in the camera position might have resulted in blurring and other errors, they also provided an opportunity to capture another appearance of the target object. We suppose that the present system can be used by non-experts, who are not nature photographers and entomologists. Therefore, our system possesses a merit for non-experts to freely change the camera’s viewpoint to capture an individual insect from one of the multiple frames.

The latest high-performance CNN-based detection and classification models require a large number of images to optimize their numerous parameters. Only a few practical and large datasets currently exist for insect detection because setting many cameras to capture numerous insects in various field backgrounds is costly (Preti et al., 2020). Although we collected hundreds of flea beetle images, merely thirty images were considered effective samples for each species. A large dataset was generated by applying data augmentation based on standard image processing such as inversion, rotation, blurring, and chroma key synthesis. Furthermore, the application of random erasing, assuming that there was an occlusion of insects in the real environment, contributed to the improvement of the accuracy in both the detection and discrimination models. The proposed image enhancement based on chroma key synthesis and random erasing was a breakthrough in the problem of research using CNNs for insect detection and identification.

Our two-stage method for tiny flea beetles may be widely applied to various insect herbivores and plants. Diverse insect herbivores, such as beetles, caterpillars, aphids and thrips, occur on *Brassica* cultivars and wild *Arabidopsis* species (Ahuja et al., 2010; Sato et al., 2019; Matsubara and Sugiura, 2021; Shimizu-Inatsugi et al., 2021). Many of these crucifer-feeding insects are as small as or larger than *Phyllotreta* beetles. For example, lepidopterous herbivores (e.g., *Plutella xylostella* and *Pieris rapae*) are recognized as a serious pest of *Brassica* crops (Ahuja et al., 2010). Last-instar larvae of them are larger than 0.5 cm and more sedentary than *Phyllotreta* beetles. In addition, *Brassica* crops are larger than our study species, *A. thaliana*. The two-stage method is therefore expected to be more feasible for large herbivores and plants than the *Phyllotreta-Arabidopsis* system. In contrast, thrips and aphids are smaller than *Phyllotreta* beetles and are likely to create a clonal colony including multiple individuals. Overlap of individuals is another issue when applying our method to the colony of small insects.

## Conclusion

To overcome the difficulty in detecting tiny insects in the field, we proposed a two-stage detection method with data augmentation. This approach improved the precision of an existing CNN-based method to help detect and identify insects in the field. To screen small insects, the YOLOv4 was used as a region proposal network. Once candidate regions were specified, another CNN-based object classification model, the EfficientNet, successfully re-identified the congeneric species from region proposals. Now that CNN-based methods have garnered much attention in biodiversity research (Norouzzadeh et al., 2018; Willi et al., 2019; Høye et al., 2021), we could combine such a state-of-the-art method to facilitate automatic identification of diverse organisms in the field.

## Acknowledgements

The authors would like to express their sincerest gratitude to H. Kuzuhara and T. Kano for the development of analytical pipelines; K. Takeda for the insect annotation; and to all members of the Shimizu group for helping with the set-up of plants in the field. This study was supported by the Japan Science and Technology Agency (JST) through the PRESTO (No. JPMJPR17Q4 to YS) and CREST (No. JPMJCR15O2 and JPMJCR16O3 to AJN and KKS) projects, the Japan Society for the Promotion Science (JSPS) KAKENHI (No. 20K15880 to YS), the Swiss National Science Foundation (No. 31003A_182318 to KKS), and the University Research Priority Project (URPP) Global Change and Biodiversity of the University of Zurich to KKS.

## Supplementary Materials

Supplementary Video S1-S3 and photographs generated by our own fieldworks are available at the GitHub repository (https://github.com/h-taki/identification-of-tiny-insects)

## References

Ahuja, I., Rohloff, J., Bones, A.M., 2010. Defence mechanisms of brassicaceae: implications for plant-insect interactions and potential for integrated pest management. a review. Agronomy for sustainable development 30, 311–348.

Almryad, A.S., Kutucu, H., 2020. Automatic identification for field butterflies by convolutional neural networks. Engineering Science and Technology, an International Journal 23, 189–195.

Bochkovskiy, A., Wang, C.Y., Liao, H.Y.M., 2020. YOLOv4: Optimal Speed and Accuracy of Object Detection. arXiv e-prints, arXiv:2004.10934arXiv:2004.10934.

Chollet, F., 2016. Xception: Deep learning with depthwise separable convolutions. Cite arxiv:1610.02357.

Dai, J., Li, Y., He, K., Sun, J., 2016. R-fcn: Object detection via region-based fully convolutional networks, in: Proceedings of the 30th International Conference on Neural Information Processing Systems, Curran Associates Inc., Red Hook, NY, USA. pp. 379–387.

Deng, L., Wang, Y., Han, Z., Yu, R., 2018. Research on insect pest image detection and recognition based on bio-inspired methods. Biosystems Engineering 169, 139–148. doi:10.1016/j.biosystemseng.2018.02.008.

Hansen, O.L., Svenning, J.C., Olsen, K., Dupont, S., Garner, B.H., Iosifidis, A., Price, B.W., Høye, T.T., 2020. Species-level image classification with convolutional neural network enables insect identification from habitus images. Ecology and Evolution 10, 737–747.

He, K., Zhang, X., Ren, S., Sun, J., 2015. Deep residual learning for image recognition. CoRR abs/1512.03385. URL: http://arxiv.org/abs/1512.03385, arXiv:1512.03385.

He, Y., Zeng, H., Fan, Y., Ji, S., Wu, J., 2019. Application of deep learning in integrated pest management: A real-time system for detection and diagnosis of oilseed rape pests. Mobile Information Systems 2019, 4570808:1–4570808:14.

Hogeweg, L., Zeegers, T., Katramados, I., Jongejans, E., 2019. Smart insect cameras. Biodiversity Information Science and Standards 3, e39241.

Howard, A.G., Zhu, M., Chen, B., Kalenichenko, D., Wang, W., Weyand, T., Andreetto, M., Adam, H., 2017. Mobilenets: Efficient convolutional neural networks for mobile vision applications. URL: http://arxiv.org/abs/1704.04861. cite arxiv:1704.04861.

Høye, T.T., Ärje, J., Bjerge, K., Hansen, O.L., Iosifidis, A., Leese, F., Mann, H.M., Meissner, K., Melvad, C., Raitoharju, J., 2021. Deep learning and computer vision will transform entomology. Proceedings of the National Academy of Sciences USA 118.

Hu, J., Shen, L., Albanie, S., Sun, G., Wu, E., 2017. Squeeze-and-excitation networks. URL: http://arxiv.org/abs/1709.01507. cite arxiv:1709.01507Comment: journal version of the CVPR 2018 paper, accepted by TPAMI.

LeCun, Y., Bengio, Y., Hinton, G., 2015. Deep learning. Nature 521, 436–444. doi:10.1038/nature14539.

Lim, S., Kim, S., Kim, D., 2017. Performance effect analysis for insect classification using convolutional neural network. 2017 7th IEEE International Conference on Control System, Computing and Engineering (ICCSCE), 210–215.

Liu, L., Ouyang, W., Wang, X., Fieguth, P., Chen, J., Liu, X., Pietikäinen, M., 2019. Deep learning for generic object detection: A survey. International Journal of Computer Vision 128, 261–318.

Liu, W., Anguelov, D., Erhan, D., Szegedy, C., Reed, S.E., Fu, C.Y., Berg, A.C., 2016. SSD: Single shot multibox detector., in: Leibe, B., Matas, J., Sebe, N., Welling, M. (Eds.), ECCV (1), Springer. pp. 21–37. doi:10.1007/978-3-319-46448-0_2.

Martineau, M., Conte, D., Raveaux, R., Arnault, I., Munier, D., Venturini, G., 2017. A survey on image-based insect classification. Pattern Recognition 65, 273–284.

Matsubara, S., Sugiura, S., 2021. A Technique for Multi-Generational Rearing of *Phaedon brassicae* (Coleoptera: Chrysomelidae). Entomological News 129, 431–434. doi:10.3157/021.129.0413.

Mayo, M., Watson, A.T., 2007. Automatic species identification of live moths. Knowledge-Based Systems 20, 195–202. doi:10.1016/j.knosys.2006.11.012. aI 2006.

Norouzzadeh, M.S., Nguyen, A., Kosmala, M., Swanson, A., Palmer, M.S., Packer, C., Clune, J., 2018. Automatically identifying, counting, and describing wild animals in camera-trap images with deep learning. Proceedings of the National Academy of Sciences USA 115, E5716–E5725.

Ovchinnikova, K., James, M.A., Mendo, T., Dawkins, M., Crall, J., Boswarva, K., 2021. Exploring the potential to use low cost imaging and an open source convolutional neural network detector to support stock assessment of the king scallop (*Pecten maximus*). Ecological Informatics 62. doi:10.1016/j.ecoinf.2021.101233.

Padilla, R., Passos, W.L., Dias, T.L.B., Netto, S.L., da Silva, E.A.B., 2021. A comparative analysis of object detection metrics with a companion open-source toolkit. Electronics 10. URL: https://www.mdpi.com/2079-9292/10/3/279, doi:10.3390/electronics10030279.

Park, J., Kim, D.I., Choi, B., Kang, W., Kwon, H.W., 2020. Classification and morphological analysis of vector mosquitoes using deep convolutional neural networks. Scientific reports 10, 1–12.

Preti, M., Verheggen, F., Angeli, S., 2020. Insect pest monitoring with camera-equipped traps: strengths and limitations. Journal of Pest Science, 1–15.

Rani, R.U., Amsini, P., 2016. Pest identification in leaf images using svm classifier. International Journal of Computational Intelligence and Informatics 6, 248–260.

Redmon, J., Divvala, S., Girshick, R., Farhadi, A., 2016. You only look once: Unified, real-time object detection, in: 2016 IEEE Conference on Computer Vision and Pattern Recognition (CVPR), pp. 779–788. doi:10.1109/CVPR.2016.91.

Redmon, J., Farhadi, A., 2018. Yolov3: An incremental improvement. ArXiv abs/1804.02767.

Ren, S., He, K., Girshick, R., Sun, J., 2017. Faster r-cnn: Towards real-time object detection with region proposal networks. IEEE Trans. Pattern Anal. Mach. Intell. 39, 1137–1149. doi:10.1109/TPAMI.2016.2577031.

Rustia, D.J.A., Lin, C.E., yung Chung, J., 2018. A real-time multi-class insect pest identification method using cascaded convolutional neural networks, in: Proceedings of 9th International Symposium on Machinery and Mechatronics for Agriculture and Biosystems Engineering, p. 67.

Samanta, R.K., Ghosh, I., 2012. Tea insect pests classification based on artificial neural networks. International Journal of Computer Engineering Science 2, 1–13.

Sato, Y., Shimizu-Inatsugi, R., Yamazaki, M., Shimizu, K.K., Nagano, A.J., 2019. Plant trichomes and a single gene *GLABRA1* contribute to insect community composition on field-grown *Arabidopsis* thaliana. BMC Plant Biology 19. doi:10.1186/s12870-019-1705-2.

Shen, Y., Zhou, H., Li, J., Jian, F., Jayas, D.S., 2018. Detection of stored-grain insects using deep learning. Computers and Electronics in Agriculture 145, 319–325. doi:10.1016/j.compag.2017.11.039.

Shimizu-Inatsugi, R., Milosavljevic, S., Shimizu, K.K., Schaepman-Strub, G., Tanoi, K., Sato, Y., 2021. Metal accumulation and its effect on leaf herbivory in an allopolyploid species *Arabidopsis kamchatica* inherited from a diploid hyperaccumulator *A. halleri*. Plant Species Biology 36, 208–217. doi:10.1111/1442-1984.12304.

Shorten, C., Khoshgoftaar, T., 2019. A survey on image data augmentation for deep learning. Journal of Big Data 6, 1–48.

Silvertown, J., Harvey, M., Greenwood, R., Dodd, M., Rosewell, J., Rebelo, T., Ansine, J., McConway, K., 2015. Crowdsourcing the identification of organisms: A case-study of iSpot. ZooKeys, 125.

Tan, M., Le, Q., 2019. EfficientNet: Rethinking model scaling for convolutional neural networks, in: Chaudhuri, K., Salakhutdinov, R. (Eds.), Proceedings of the 36th International Conference on Machine Learning, PMLR. pp. 6105–6114.

Valan, M., Makonyi, K., Maki, A., Vondráček, D., Ronquist, F., 2019. Automated taxonomic identification of insects with expert-level accuracy using effective feature transfer from convolutional networks. Systematic Biology 68, 876–895.

Van Horn, G., Mac Aodha, O., Song, Y., Cui, Y., Sun, C., Shepard, A., Adam, H., Perona, P., Belongie, S., 2018. The iNaturalist species classification and detection dataset, in: Proceedings of the IEEE conference on computer vision and pattern recognition, pp. 8769–8778.

Venugoban, K., Ramanan, A., 2014. Image classification of paddy field insect pests using gradient-based features. International Journal of Machine Learning and Computing, 1–5.

Wang, J., Lin, C., Ji, L., Liang, A., 2012. A new automatic identification system of insect images at the order level. Knowledge-Based Systems 33, 102–110. doi:10.1016/j.knosys.2012.03.014.

Willi, M., Pitman, R.T., Cardoso, A.W., Locke, C., Swanson, A., Boyer, A., Veldthuis, M., Fortson, L., 2019. Identifying animal species in camera trap images using deep learning and citizen science. Methods in Ecology and Evolution 10, 80–91.

Xia, D., Chen, P., Wang, B., Zhang, J., Xie, C., 2018. Insect detection and classification based on an improved convolutional neural network. Sensors 18. doi:10.3390/s18124169.

Xie, C., Wang, R., Zhang, J., Chen, P., Dong, W., Li, R., Chen, T., Chen, H., 2018. Multi-level learning features for automatic classification of field crop pests. Computers and Electronics in Agriculture 152, 233–241. doi:10.1016/j.compag.2018.07.014.

Xie, C., Zhang, J., Li, R., Li, J., Hong, P., Xia, J., Chen, P., 2015. Automatic classification for field crop insects via multiple-task sparse representation and multiple-kernel learning. Computers and Electronics in Agriculture 119, 123–132. doi:10.1016/j.compag.2015.10.015.

Yalcin, H., 2015. Vision based automatic inspection of insects in pheromone traps, in: 2015 Fourth International Conference on Agro-Geoinformatics (Agro-geoinformatics), pp. 333–338. doi:10.1109/Agro-Geoinformatics.2015.7248113.

Yamashita, R., Nishio, M., Do, R., Togashi, K., 2018. Convolutional neural networks: an overview and application in radiology. Insights into Imaging 9, 611–629. doi:10.1007/s13244-018-0639-9.

Zhong, Z., Zheng, L., Kang, G., Li, S., Yang, Y., 2020. Random erasing data augmentation, in: The Thirty-Fourth AAAI Conference on Artificial Intelligence, AAAI 2020, The Thirty-Second Innovative Applications of Artificial Intelligence Conference, IAAI 2020, The Tenth AAAI Symposium on Educational Advances in Artificial Intelligence, EAAI 2020, New York, NY, USA, February 7-12, 2020, AAAI Press. pp. 13001–13008. doi:10.1609/aaai.v34i07.7000.

